# Mitochondrial NADK2-dependent NADPH controls Tau oligomer uptake in human neurons

**DOI:** 10.1101/2024.10.31.621392

**Authors:** Evelyn Pardo, Taylor Kim, Horst Wallrabe, Kristine E. Zengeler, Vijay Kumar Sagar, Garnett Mingledorff, Xuehan Sun, Ammasi Periasamy, John R. Lukens, George S. Bloom, Andrés Norambuena

## Abstract

Alterations in NADH and NADPH metabolism are associated with aging, cancer, and Alzheimer’s Disease. Using 2P-FLIM imaging of the mitochondrial NAD(P)H in live human neurons and PS19 mouse brains, we show that tau oligomers (TauO) upregulate the mitochondrial *de novo* NADPH synthesis through NADK2. This process controls LRP1-mediated internalization of TauO, setting a vicious cycle for further TauO internalization. Thus, mitochondrial NADK2-dependent NADPH controls a key step in TauO toxicity.

---

Neuronal uptake of pathological forms of tau^1–5^, mitochondrial dysfunction^6^ and brain hypometabolism^7,8^ are key features in Alzheimer’s disease (AD) pathogenesis; how these processes are mechanistically connected remains elusive. Mitochondria are key players in energy metabolism, redox regulation, and biosynthetic pathways^9^; these functions directly depend on the coenzymes contained within the organelle. Two important coenzymes are the pyridine nucleotide NADH and its phosphorylated form NADPH, which regulate oxidative phosphorylation and provide reducing power for biosynthetic pathways, respectively^10^ (Fig. 1a). There are several cellular sources of NADPH^10^, but its *de novo* synthesis in mitochondria requires NAD kinase, NADK2^11,12^, whose role in AD is unknown.

**Figure 1.**
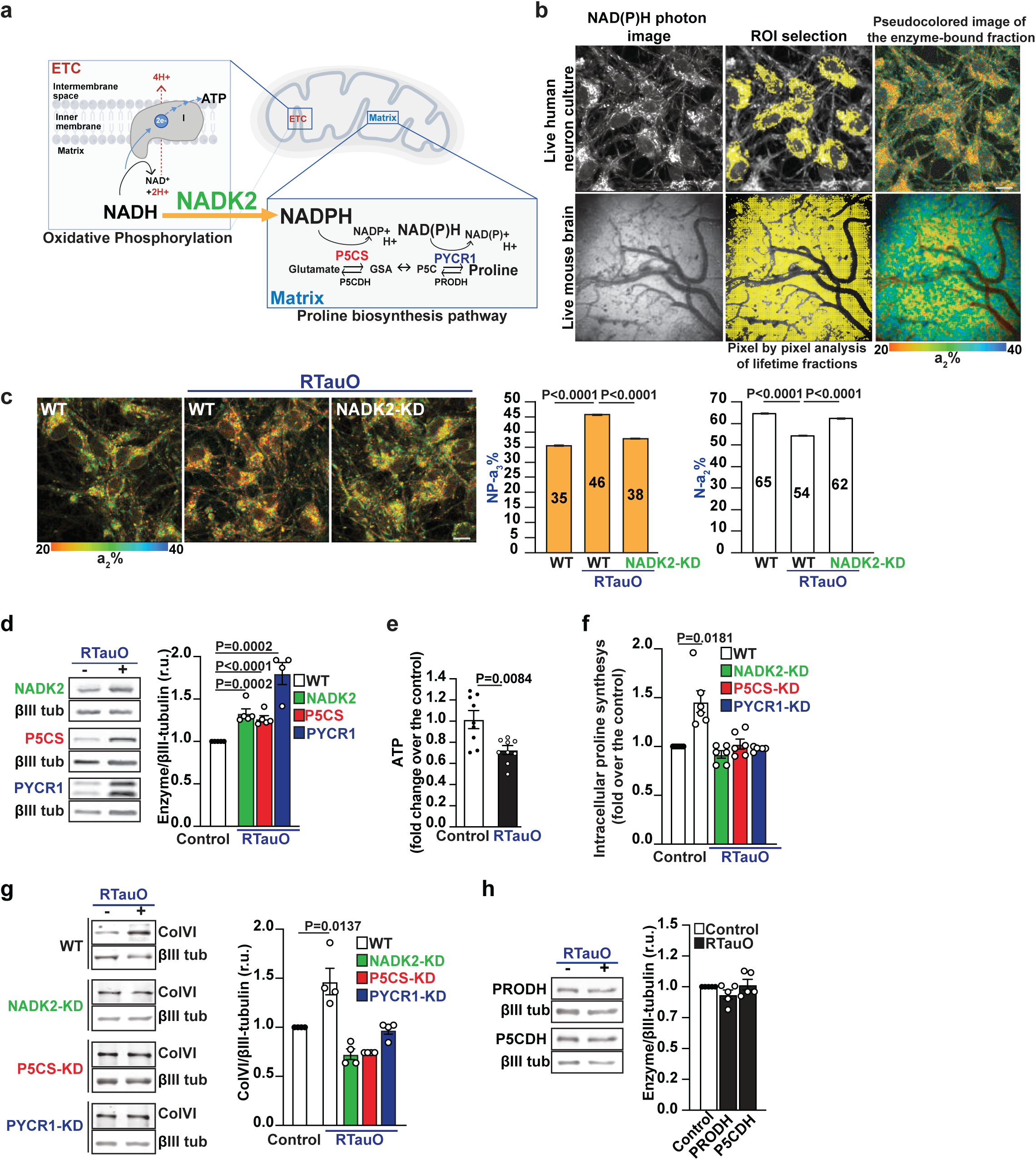
TauO increases the expression of mitochondrial NADPH and proline synthesis pathway in a NADK2-dependent manner. **a.** A simplified cartoon depicting the roles of NADH and NADPH [NAD(P)H] in mitochondria. NAD(P)H coenzymes act as electron carriers in metabolic reactions. NADH mainly drives ATP synthesis through mitochondrial oxidative phosphorylation, and NADPH mostly supports reductive biosynthesis (e.g., proline biosynthesis). The mitochondrial *de novo* synthesis of NADPH depends on the NADK2. **b.** Two-photon fluorescence lifetime microscopy (2P-FLIM) detects the total auto-fluorescent lifetime of the mitochondrial enriched fraction of NADH and NADPH bound to enzyme partners, both in live neuronal cultures and live mouse brains. This assay allows label-free imaging of mitochondrial activity in live specimens by tracking the differential lifetime of free coenzymes Ꚍ1 (∼400 ps) and their fraction a_1_%, enzyme-bound NADH-Ꚍ2 (∼1500-4400 ps) and their fraction a_2_% (N-a_2_%) and enzyme-bound NADPH-Ꚍ3 (>4400 ps) and their fraction a_3_% (NP-a_3_%). As described in M&M, the images (top panel, live human neuronal culture; bottom panel mouse brains) represent the analysis sequence, starting with collecting the photon images, continuing with ROI selection, and ultimately extracting the N-a2% and NP-a3% fractions. **c.** RTauO stimulates the NADK2-dependent NADPH in human neurons. Representative 2P-FLIM images of WT or NADK2-KD neurons treated with a vehicle (control) or 100 nM of bacterially produced recombinant 2N4R human tau oligomers (RTauO) for 24 hours. 2P-FLIM was used to monitor changes in N-a_2_% and NP-a_3_%. RTauO-mediated increase in NADPH was significantly reduced in NADK2-KD neurons. Scale bar, 20μm. As more than 5,000 ROIs were obtained per image, the average ROI for each field of view was calculated. Statistical analyses were performed using a two-tailed unpaired student’s t-test. Error bars represent +/-s.e.m. (n=3). **d.** WT human neuron cultures were treated with a vehicle (control) or 100 nM of RTauO for 24 hours. NADK2 (n=5), P5CS (n=4), and PYCR1 (n=5) levels were measured by western blot (WB) and normalized to the expression level of βIII tubulin contained in the same samples. Statistical analyses were performed using a two-tailed unpaired student’s t-test. Error bars represent +/- s.e.m. **e.** ATP levels were measured in WT human neurons treated with a vehicle (control) or 100 nM of RTauO for 24 hours using the ATPlite 1step Luminescence Assay System (Perkin-Elmer). The data were collected from three independent assays, each containing three replicates per experimental condition. Error bars represent +/- s.e.m. **f.** Intracellular proline levels were measured using the acid-ninhydrin assay in either WT, NADK2-KD, P5CS-KD, or PYCR1-KD neurons treated with vehicle (control) or 100 nM of RTauOs for 24 hours. Statistical analyses were performed using a two-tailed unpaired student’s t-test. Error bars represent +/- s.e.m. (n=6). **g.** Collagen-VIa (ColVI), another read-out for proline synthesis pathway, was analyzed in either WT, NADK2-KD, P5CS-KD, or PYCR1-KD neurons treated with vehicle (control) or 100 nM of RTauOs for 24 hours. ColVI expression detected by WB was normalized to the expression of βIII tubulin contained in the same samples. (n=4). **h.** TauO does not change the expression of enzymes required for proline catabolism. WT human neuron cultures were treated with a vehicle (control) or 100 nM of RTauO for 24 hours. PRODH and P5CDH levels were measured by WB and normalized to the expression of βIII tubulin contained in the same samples. Statistical analyses were performed using a two-tailed unpaired student’s t-test. Error bars represent +/- s.e.m. (n=5).

To study the effect of tau oligomers (TauO) on mitochondrial metabolism in live neurons and mouse cortex, we use label-free, 2-photon fluorescence lifetime microscopy (2P-FLIM) to image NAD(P)H lifetimes^13–15^. Tracked changes in the coenzyme’s fluorescent lifetime fractions of biochemically active, enzyme-bound NADH-a_2_% (N-a_2_%) or NADPH-a_3_% (NP-a_3_%) act as a proxy for mitochondrial respiration for the former^13^, and activation of biosynthetic pathways for the latter (Fig. 1a, b)^11,12^. While spectrally NADPH and NADH are alike, their fluorescence lifetimes and fractions are discriminated by 2P-FLIM^16^, allowing measurement of the relative contribution of each coenzyme in live specimens. Human neurons derived from ReNCell™ MV human neuronal progenitor cell line^13,15^ (hereafter human neurons) were vehicle-treated (control) or exposed for 24 h to 100 nM of bacterially produced recombinant 2N4R human tau oligomers (RTauO)^17^ (Extended Data Fig. 1a, b). Under control conditions, NP-a_3_% contributed ∼35% of the fraction, while N-a_2_% accounted for ∼65% (Fig. 1c). Interestingly, RTauO treatment led to a ∼31% increase in NP-a_3_% and the corresponding decrease in N-a_2_% in WT neurons (Fig. 1c). Additionally, RTauO also upregulated the expression of NADK2 in human neurons by ∼30% (Fig. 1d), which agrees with the fact that NADPH availability correlate with kinase abundance^18^. Reciprocally, RTauO-mediated increase in NP-a_3_% was reduced by ∼22.5% in NADK2 knocked-down neurons (Fig. 1c and Extended Data Fig. 2d), further indicating the specificity of our 2P-FLIM assay. Additionally, RTauO-mediated imbalance in mitochondrial NADH and NADPH content lead to a 20% decrease in total ATP, likely because of lower availability of NADH to feed the Complex I of the electron transport chain (Fig. 1e).

It has been recently reported that NADK2-dependent NADPH regulates mitochondrial proline biosynthesis (Fig. 1a)^11,12^; this suggests that TauO could activates the mitochondrial proline synthesis pathway. Mitochondrial-dependent biosynthesis of proline is catalyzed by specific enzymes. First, P5C synthase (P5CS) converts glutamate to glutamic-γ-semialdehyde, which spontaneously cycles to Δ1-pyrroline-5-carboxylate (P5C). This step is accompanied by the transfer of a reducing power from NADPH. Then, P5C is catalyzed to L-proline by P5C-reductase (PYCR1), followed by the transfer reduction potential from NADH or NADPH^19^. These steps affect the mitochondrial content of NADH and NADPH and the associated biological processes regulated by them^20^ (Fig. 1a). RTauO not only increased NP-a_3_% and NADK2 (Fig. 1c, d) but also upregulated the expression of P5CS and PYCR1 (Fig. 1d) and increased the neuronal levels of proline and collagen-VIa (ColVI), which is a well-known product of the proline synthesis activation^21^ and is overexpress in AD models^22^ (Fig. 1f, g). These outcomes were not observed in NADK2, P5CS, or PYCR1 knockdown human neurons (Fig. 1f, g, and Extended Data Fig. 2d-f). In addition, RTauO did not affect the expression levels of PRODH and P5CDH, which regulate proline catabolism (Fig. 1h)^11,12^. Importantly, RTauO did not change the expression of NADK1, the kinase responsible for cytosolic *de novo* synthesis of NADPH (Extended Data Fig. 3a). Consequently, RTauO-mediated upregulation of NADK2, proline, and ColVI synthesis was unaffected in NADK1 knocked down neurons (Extended Data Fig. 3b-d, f).

NADK2 plays a role in cell proliferation and cancer^11,12^, and mutations in this enzyme have major neurological consequences associated with early death in humans^23,24^; however, the mechanisms involved are unknown. To better understand the molecular pathways regulated by NADK2, we performed quantitative protein mass spectrometry (MS) in NADK2-knockout (NADK2-KO; Extended Data Fig. 2a) and WT human neurons. The analysis identified 371 proteins whose expression was significantly affected by NADK2 expression (Table 1; P<0.01299), including 188 proteins whose expression were significantly reduced in NADK2-KO neurons (Fig. 2a). A key step in TauO toxicity is its endocytic uptake by receiving cells; accordingly, we found that 17 proteins (4.5%) were involved in the regulation of membrane trafficking. Remarkably, we found that the expression of the low-density lipoprotein receptor-related protein 1 (LRP1), a major tau receptor at the plasma membrane^5^, was reduced by 50% in NADK2-KO neurons (Fig. 2a). No other proteins known to be involved in TauO internalization were identified. Consistent with the observation that RTauO upregulates both NP-a_3_% (Fig. 1c) and NADK2 expression (Fig. 1d) along with proline and ColVI biosynthesis (Fig. 1f, g), RTauO treatment also increased LRP1 expression by more than 50% in WT but not in human neurons deficient in either NADK2, P5CS or PYCR1 (Fig. 2b and Extended Data Fig. 2a-c). In agreement with the role of LRP1 in regulating TauO uptake, RTauO-mediated increase in mitochondrial NP-a_3_%, proline, and ColVI biosynthesis were obliterated in LRP1 knocked down neurons (Fig. 2c-e and Extended Data Fig. 2g). Thus, our observations suggest that TauO orchestrate mitochondrial functioning and membrane trafficking to facilitate their uptake.

**Figure 2:**
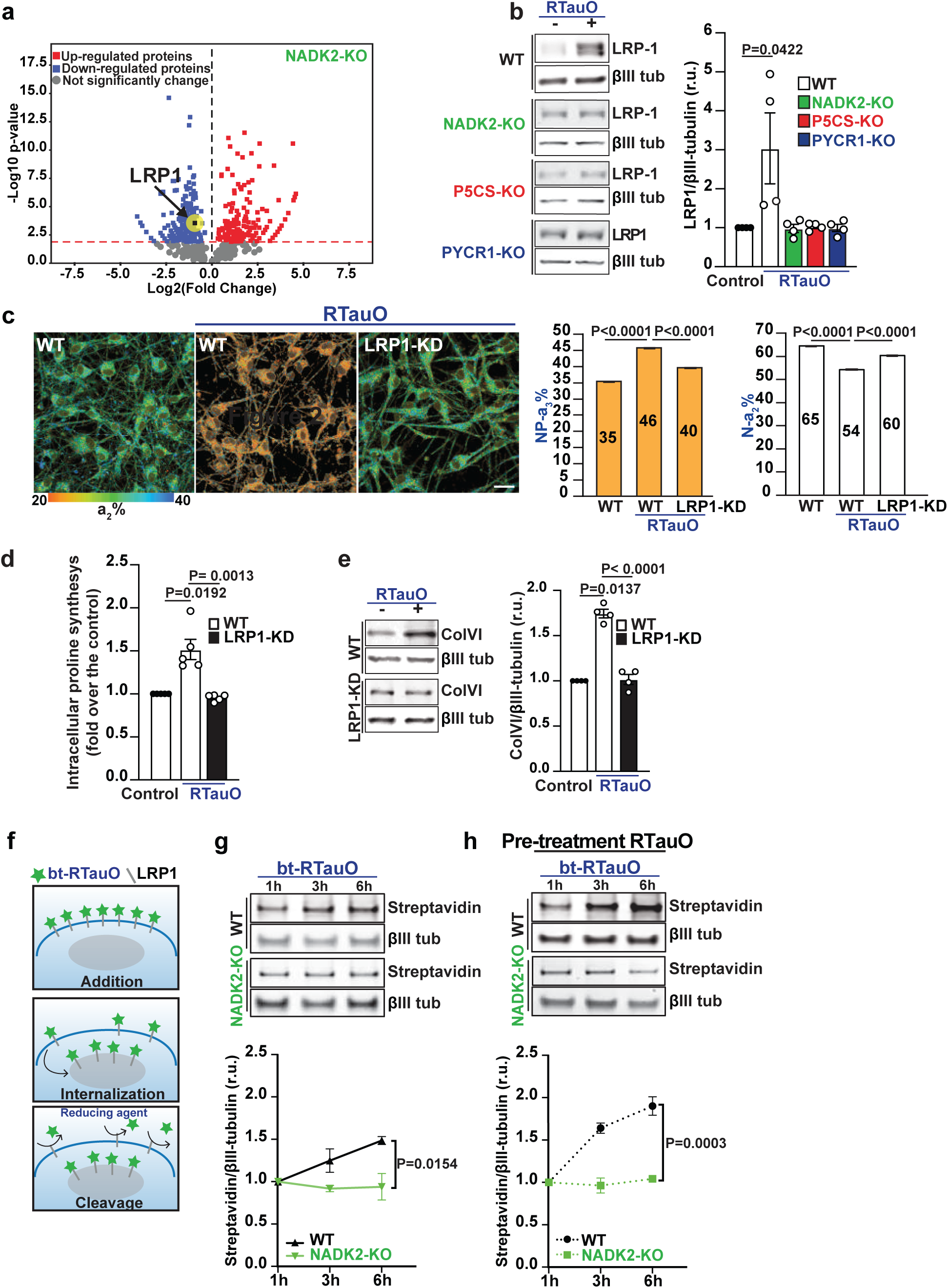
NADK2 controls LRP1 expression and TauO endocytosis. **a.** Proteomic analysis from postnuclear membrane fractions obtained from WT or NADK2-KO human neurons measured by LC/MS/MS. Significantly changed proteins were determined using Scaffold software (v5.3) (Fisher’s Exact Test, p<0.01299, Benjamini-Hochberg). Downregulated (blue) and upregulated (red) proteins in NADK2-KO neurons are represented in the volcano plot (Detailed in Table 1). **b.** LRP1 expression was analyzed by WB of protein samples from human neurons treated for 24 hours with vehicle (control) or 100 nM RTauO. The results were compared to either NADK2-KO, P5CS-KO, or PYCR1-KO neurons. LRP1 expression level was normalized to the expression level of βIII tubulin contained in the same samples. Statistical analyses were performed using a two-tailed unpaired student’s t-test. Error bars represent +/- s.e.m. (n=4). **c.** LRP1 controls TauO-mediated increment of mitochondrial NP-a_3_%, intracellular proline, and collagen synthesis. Representative 2P-FLIM images of WT or LRP1-KD neurons were treated with vehicle (control) or 100 nM of RTauO for 24 hours, and 2P-FLIM imaging was used to analyze N-a_2_% and NP-a_3_%. Scale bars, 10μm. Because thousands of ROIs were obtained per image, the average ROI on each field of view was calculated. Error bars represent +/- s.e.m. n=3. **d.** Intracellular proline levels were measured using the acid-ninhydrin assay in WT human neurons treated with a vehicle (control) or 100 nM of RTauOs for 24 hours, and the results were compared to LRP1-KD neurons. Statistical analyses were performed using a two-tailed unpaired student’s t-test. Error bars represent +/- s.e.m. (n=5)**. e.** In a similar experiment, ColVI synthesis was monitored by WB and normalized to the expression level of βIII tubulin contained in the same samples. (n=4). Statistical analyses were performed using a two-tailed unpaired student’s t-test. Error bars represent +/- s.e.m. **f.** Scheme showing the internalization protocol of biotin-tagged RTauO (bt-RTauO). **g.** NADK2 controls TauO uptake by neurons. 300 nM of bt-RTauO were added to either WT or NADK2-KO neuron cultures at 37°C for the indicated time points. Then, the internalized (DTT-resistant) bt-RTauO was monitored by WB using iRDye800-labeled streptavidin (LICOR). While the internalization of bt-RTauO in WT neurons increased by ∼50% in a 6 h time course, the corresponding incorporation in NADK2-KO neurons remains steady. Line graphs show the quantification obtained using 3 independent experiments and analyzed using a two-tailed unpaired student’s t-test. Internalized bt-TauO was normalized to the expression level of βIII tubulin contained in the same samples. Error bars represent +/- s.e.m. **h.** TauO stimulated its own uptake by a NADK2-dependent mechanism. WT and NADK2-KO neurons were treated with 100 nM RTauO for 24 hours before performing the bt-RTauO uptake assay, as shown in **g**. Internalized bt-TauO was normalized to the expression level of βIII tubulin contained in the same samples. Line graphs show the quantification obtained using 3 independent experiments and analyzed using a two-tailed unpaired student’s t-test. Error bars represent +/- s.e.m. n=3.

How TauO impacts cellular functions is a matter of intense investigation; a critical step is their uptake via endocytosis, a process mediated in part by LRP1^5^. As TauO controls the expression of LRP1 in a NADK2-dependent manner (Fig. 2b), we asked whether NADK2 regulates the uptake of TauO by neurons. RTauO were labeled with EZ-Link™-Sulfo-NHS-SS-Biotin, which allows reversible labeling of primary amines in the proteins. Human neurons were exposed to bt-RTauO for different time periods. Before sample harvest, biotin from non-internalized bt-RTauO was washed off by a buffer containing DTT, as internalized bt-RTauO were protected from the reduction (Fig. 2f). While the incorporation of bt-RTauO showed a steady uptake in WT neurons in a 6-hour time course, its uptake in NADK2-KO cells was reduced by ∼50% (Fig. 2g). Knocking down NADK1 expression did not affect bt-RTauO uptake (Extended Data Fig. 3e). Interestingly, upregulating the expression of LRP1 (by pretreating neurons with 100 nM of RTauO; Fig. 2b) for 24 h prior the internalization experiment, further increased the uptake of bt-RTauO in WT neurons by 25%, but not in NADK2-KO neurons (Fig. 2h). Thus, TauO initiates a vicious cycle of uptake by upregulating the mitochondrial NADK2-dependent NAPDH.

To understand the relevance of these observations to human AD, we treated human neurons with TauO purified from human AD brain (<u>b</u>rain-<u>d</u>erived <u>TauO</u>; BDTauO). Importantly, BDTauO treatment also increased the expression of NADK2 and a corresponding NADK2-dependent increase of NP-a_3_% (Fig. 3a, b). BDTauO also upregulated LRP1 overexpression by 50% in a NADK2-dependent manner (Fig. 3c). Additionally, NADK2 and PYCR1 expression was upregulated in brain cell extracts obtained from deceased human AD patients compared to aged-matched, cognitively normal donors (Fig. 3d).

**Figure 3:**
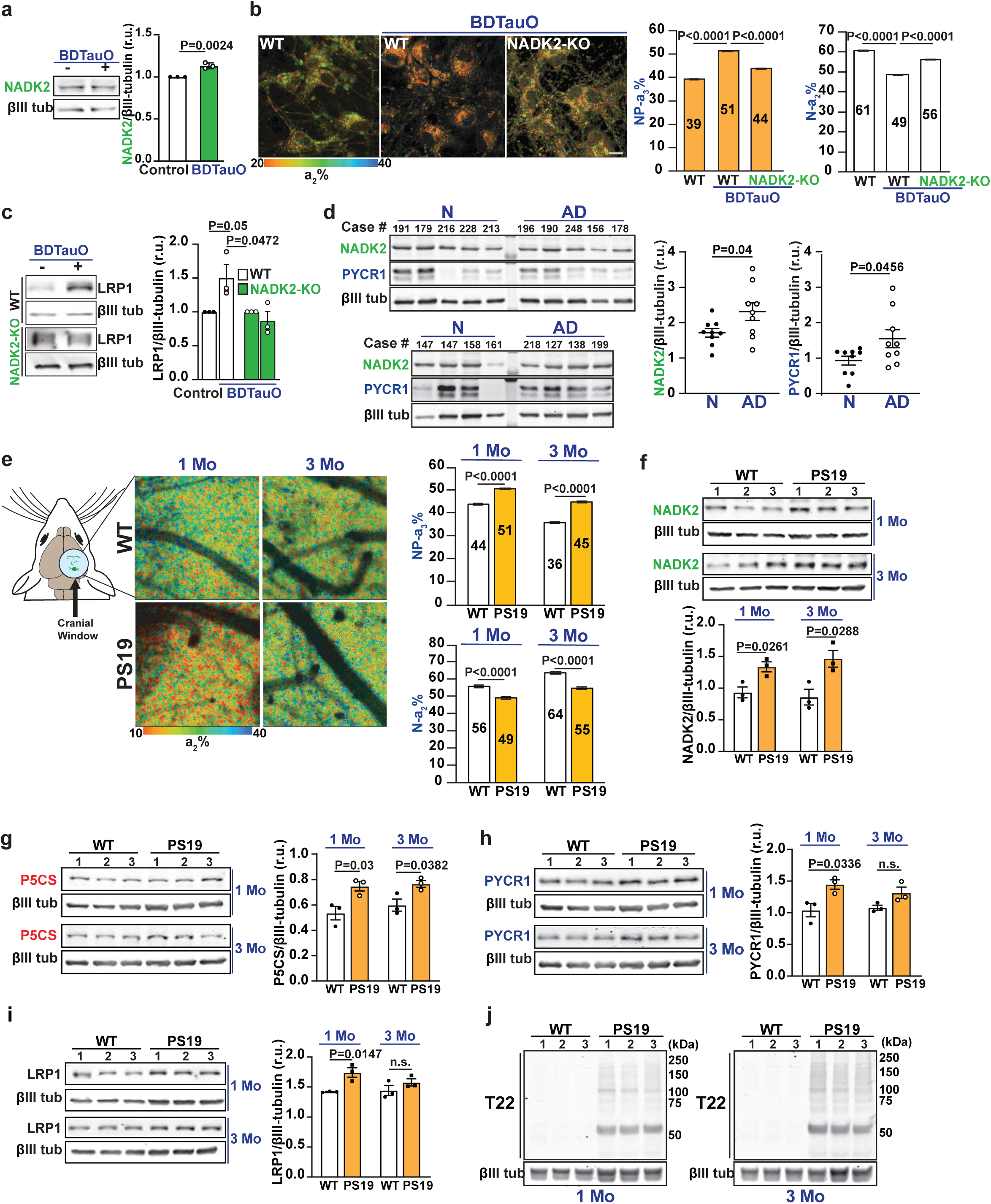
Human AD brain-derived tau oligomers increase NADK2-dependent NADPH in neuron cultures and live mouse brains. Human AD brain-derived TauO stimulates the NADK2-dependent NADPH. **a.** WT human neuron cultures were treated with a vehicle (control) or 100 nM of tau oligomers purified from human AD brain (<u>b</u>rain-<u>d</u>erived <u>TauO</u>; BDTauO) for 24 hours. NADK2 levels were measured by WB (n=3) and normalized to the expression level of βIII tubulin contained in the same samples. Statistical analyses were performed using a two-tailed unpaired student’s t-test. Error bars represent +/- s.e.m. **b.** Representative 2P-FLIM images of WT or NADK2-KO neurons treated with a vehicle (control) or 100 nM of BDTauO for 24 hours. 2P-FLIM was used to monitor changes in N-a_2_% and NP-a_3_%. BDTauO-mediated increase in NADPH was significantly reduced in NADK2-KO neurons. Scale bars, 20μm. Because thousands of ROIs were obtained per image, the average ROI for each field of view was calculated. Statistical analyses were performed using a two-tailed unpaired student’s t-test. Error bars represent +/- s.e.m. (n=3, two different batches of BDTauO from two different AD patients). **c.** LRP1 expression levels in WT neurons were analyzed as described above and compared to NADK2-KO neurons (n=3). LRP1 expression was normalized to the expression level of βIII tubulin contained in the same samples. Statistical analyses were performed using a two-tailed unpaired student’s t-test. Error bars represent +/- s.e.m. **d.** NADK2 and PYCR1 expression were analyzed in human brain cell extracts obtained from deceased patients affected by AD and compared to cognitively normal controls (same samples described by us^1^). Samples were analyzed by western blots, and the expression of either NADK2 or PYCR1 was normalized to the expression level of βIII tubulin contained in the same samples. Statistical analyses were performed using a two-tailed unpaired student’s t-test. Error bars represent +/- s.e.m. **e.** Representative 2P-FLIM imaging in WT and presymptomatic PS19 mouse cortex through an open-skull window. An increase in NP-a_3_% was observed in the PS19 mouse cortex, likely indicating upregulation of the mitochondrial NADK2-dependent NADPH. Statistical analyses were performed using a two-tailed unpaired student’s t-test. 3 animals/age/groups/genetic backgrounds were used for recording N-a_2_% and NP-a_3_%. Error bars represent +/- s.e.m. **f.** NADK2 is upregulated in presymptomatic PS19 mice. Cortical protein samples from WT or PS19 brains were separated by SDS-PAGE, followed by WB of the indicated proteins. The expression levels of NADK2 were normalized to the one of βIII tubulin detected on each sample. Bar graphs show the quantification obtained using 3 mice/group/genetic background and analyzed using a two-tailed unpaired student’s t-test. Increased NADK2 and NADPH in the cortex of presymptomatic PS19 mice correlated with the expression of tau proteoforms. Protein samples from WT and PS19 mice cortex were analyzed by WB using antibodies against P5CS (**g**), PYCR1 (**h**), LRP1 (**i**), and T22, which detects tau proteoforms (**j**). The expression levels of these proteins were normalized to the expression level of βIII tubulin contained in the same samples. The bar graph shows the quantification obtained using 3 mice/group/genetic background and analyzed using a two-tailed unpaired student’s t-test. Error bars represent +/ - s.e.m.

The generation of human-derived neurons (iNeurons) through direct conversion of dermal fibroblasts has facilitated the exploration of molecular mechanisms of age-related human diseases such as AD^25^. Here, iNeurons obtained from AD patients (Extended Data Fig. 4a) displayed higher levels of NP-a_3_% compared to iNeurons from cognitively normal controls (Extended Data Fig. 4b). Like our observations with RTauO and BDTauO (Fig. 1d; Fig. 3a), higher NP-a_3_% values in AD iNeurons also coincided with an upregulation in the expression of NADK2, P5CS, PYCR1and LRP1 (Extended Data Fig. 4c, d) and, strikingly, with the detection of T22 antibody^26^-positive tau proteoforms (Extended Data Fig. 4e). Thus, upregulation of the NADK2-NADPH and proline synthesis pathway might represent a novel human AD phenotype.

Pathological tau (p181 and p217) has been detected in CSF and plasma of human patients >10 years before the onset of cognitive decline^27^, suggesting that soluble forms of tau could impair mitochondrial metabolism *in vivo* years before disease onset. We took advantage of PS19 mice, which naturally produce and accumulate TauO in the brain months before synaptic abnormalities and cognitive deficits occur^28^. We recorded the energy metabolic state in the cortex of the PS19 live mouse brain at 1 and 3 months of age using 2P-FLIM, which allows measuring mitochondria activity *in vivo* by tracking N-a_2_% and NP-a_3_% through an open-skull window^15^. Compared to WT littermates, PS19 mice showed a time-dependent increase of NP-a_3_% by 16% at 1 month age and 25 % at 3 months of age (Fig. 3e). The rise of NP-a_3_% in presymptomatic PS19 mice paralleled the upregulation of NADK2, P5CS, PYCR1 and LRP1 expression in the cortex of these animals (Fig. 3f- i) and the early detection of T22 antibody-positive tau proteoforms (Fig. 3j). Altogether, these results show that TauO upregulates the expression of LRP1 *in vivo* by a NADK2-dependent mechanism, highlighting that TauO manifests its toxicity through dysregulation of mitochondrial NADK2.

NADH and NADPH coenzymes function as cofactors and/or substrates for numerous enzymes to maintain cellular redox balance, energy metabolism, and other biological processes^29^. Prolonged disequilibrium of NADH/NADPH is associated with diseases such as cancer, aging, and neurodegenerative disorders^10^. Here, we provide evidence that TauO controls its own uptake by upregulating the mitochondrial NADK2-dependent NADPH and the expression of LRP1 (Fig. 4) likely at presymptomatic stages of AD. Since LRP1 is a main regulator of TauO uptake and spread^5^, dysregulation of NADPH balance could be a very early contributor to human AD initiation.

**Figure 4:**
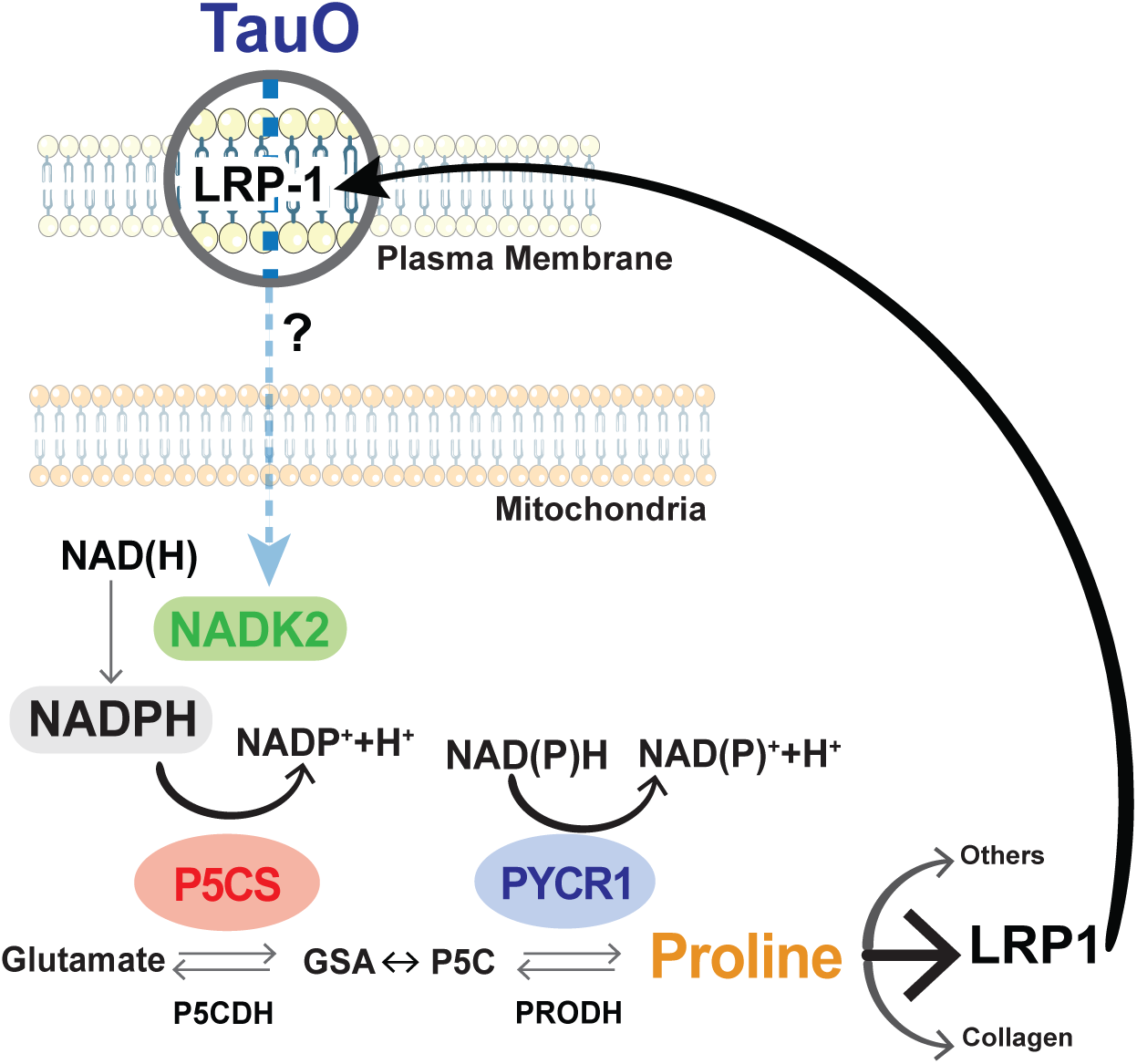
Working Model Mitochondrial NADK2-dependent NADPH controls Tau oligomer uptake in human neurons. Neurons exposed to soluble and oligomeric forms of TauO upregulate the mitochondrial *de novo* NADPH synthesis by upregulating NADK2. Importantly, upregulation of NADPH was observed in PS19 live mouse brain (which naturally produce human TauO) months before cognitive decline onset, and in “inducible AD neurons” obtained from direct conversion of human dermal fibroblasts. Activation of this NADK2-dependent biochemical pathway then controls the expression of the LRP1, a main TauO receptor in the neuronal plasma membrane. This vicious cycle, whereby TauO orchestrates mitochondrial NADPH metabolism and membrane trafficking to stimulate their own uptake, implies that TauO-mediated dysregulation of NADK2 is one of the earliest molecular missteps in AD pathogenesis.

## Acknowledgments

We would like to thank Dr. Anthony Spano for providing the βIII-tubulin antibody (Tuj1) and Dr. Rakez Kayed for his generous gift of human brain-derived Tau oligomers. We also thank the W.M. Keck Biomedical Mass Spectrometry Laboratory (funded by a grant from the University of Virginia’s School of Medicine), and especially Dr. Dilza Silva for helpful discussions.

## Consent Statement

Informed consent was obtained from all subjects before sample collection. Receipt of human tissues was granted through University of Virginia’s Institutional Review Board for Health Sciences Research.

## Author contributions

EP and AN conceived, designed, initiated, and led the entire study. EP performed all the studies of neurons in culture, took care of the animal colony, performed genotyping, and along with AN and VS coordinated live mouse experiments. EP generated CRISPR/CAS9 cells and wrote the first draft of the paper, and, along with AN, served as the main editor of the original text. VS performed 2P-FLIM experiments in the live mouse brain. HW and VS analyzed 2P-FLIM experiments in the live cells and mouse brains. TK provided technical support throughout the study. AP provided extensive technical support for 2P-FLIM. XS generated the initial TauO preparation and prepared samples for TauO negative staining and Atomic Force microscopy. GM performed the culture of iNeurons. KZ and JL provided extensive technical and intellectual support for in vivo experiments. GSB analyzed data and provided intellectual feedback throughout the study. All authors read, edited, and approved the submission of the manuscript.

## Materials and Methods

### Cell Culture and materials

Human embryonic kidney cells (Lenti-X 293T Cell line from Takara. Cat no. 632180) were grown in DMEM/F12 media (GIBCO) supplemented with HyClone cosmic calf serum (GE Healthcare) and 50 µg/ml gentamycin (GIBCO).

As described previously ^1–3^, human neurons were differentiated from ReNCell VM neuronal precursor cells (EMD Millipore). For differentiation, the cells were plated into 12 wells plates or 35 mm glass-bottom dishes with DMEM/F12 differentiation media (GIBCO) supplemented with 2 µg/ml heparin (Science Cell), 2% (v/v) B27 neural supplement (GIBCO), and 50 µg/ml gentamycin (GIBCO) without growth factors. One-half volume of the differentiation media was changed every three days until the cultures were used for experiments.

### Preparation of iNeurons

Direct Conversion of Adult Human Fibroblasts into iNeurons (iN). Primary human dermal fibroblasts were obtained from the Coriell Institute Cell Repository and patients enrolled in study HSR210075 at the University of Virginia. Protocols were previously approved by the UVA Internal Review Board for Health Sciences Research, and informed consent was obtained from all subjects. Fibroblasts were cultured in MEM (Gibco #11095-080) containing 15% tetracycline-free fetal bovine serum (VWR 76308-984), 1% NEAA (Gibco #11140050), 1 mM Sodium Pyruvate (Gibco #11360070) and transduced with lentivirus expressing the transcription factors Ngn2:2A and Ascl1. Transduced fibroblasts were expanded with 1 µg/ml puromycin (Gibco A1113803). Following at least four to five passages post-transduction, cells were considered ’iN-ready.’ They were trypsinized and pooled into high densities (∼140,000–500,000 cells per cm^²^ in 2D & 3D culture, respectively) and plated in optically clear 24-well plates (Ibidi USA #82426). 3D cultures were resuspended in a media/Matrigel mix (5mg/ml final concentration: Corning #356231) and allowed to polymerize at 37°C for 40 min before adding additional media. The following day, the medium was changed to neuron conversion medium (NC) based on a 1:1 mix of DMEM: F12 (Gibco #11320033) and Neurobasal-A (Gibco #10888022) for three to five weeks. NC contains the following supplements: 1x N2 (Gibco #17502048), 1x B27 (Gibco #17504044), 2 µg/ml doxycycline (Sigma #D9891), 1 µg/ml Laminin (Thermo-Fisher #23017015), 400 µg/ml dibutyryl-cyclic-AMP (Sigma #D0627), 150 ng/ml human recombinant Noggin (PeproTech #120-10C), 0.5 µM LDN-193189 (Cayman Chemicals #19396) 0.5 µM A83-1 (Santa Cruz #sc-203791), 3 µM CHIR99021 (Cayman Chemicals #13122), 5 µM Forskolin (BioGems #6652995) and 10 µM SB-431542 (Cayman Chemicals #13031). The medium was changed every two to three days. Once neuronal morphology was detected (weeks 2-3), 10% Astrocyte Conditioned Medium was supplemented (ScienCell #1811) to the NC medium.

### Mouse Strain

The PS19 (**B6.Cg-Tg(Prnp-MAPT*P301S)PS19Vle/J**) tauopathy mouse model was purchased from The Jackson Laboratory (Strain #024841). Animals were housed in a pathogen-free barrier facility with a 12-hour light/12-hour dark cycle and ad libitum access to food and water under a protocol approved by the IACUC of the University of Virginia. All pups were genotyped by PCR, following The Jackson Laboratory’s recommendations. All experiments were performed in males for PS19 and WT littermates.

### Lentivirus production and transduction

Lentiviral particles for shNADK2, shP5CS, shPYCR1, and shLRP1 knockdowns were prepared as follows. The expression plasmids, pLKO.1 (Mission shRNA library from Sigma-Aldrich, see below), and the packaging vectors, pSPAX2 and pMD2.G (Addgene plasmids #12260 and #12259, respectively) were transfected using Lipofectamine 3000 (Thermo-Fisher) into HEK293T cells grown in 15 cm Petri dishes to ∼70% confluence in DMEM (GIBCO) supplemented with 10% HyClone cosmic calf serum. Each transfection was performed with 15 µg total DNA at a 50% expression vector, 37.5% pSPAX plasmid, and 12.5% pMD2G plasmid. The lentivirus-conditioned medium was collected 24 and 48 hours after the start of transfection. For expression of Ngn2:2A and Ascl1 in dermal fibroblasts, Hek 293 cells were co-transfected with pLVX-UbC-rtTA-Ngn2:2A:Ascl1 (Addgene #127289) and Lenti-X Packaging Single Shots (VSV-G; Takara #631275) following the manufacturer recommendations. Lentiviral particles were concentrated in a Beckman Coulter Optima XE-90 ultracentrifuge for 2 hours at 23,000 rpm (90,353 x g_av_) at 4° C in an SW32Ti rotor, resuspended in 400 µl Neurobasal medium and stored at −80° C in 40 µl aliquots. Cultured neurons were transduced for 72-96 hours before the experiments were performed.

### Human Brain Cell Extracts

The Institutional Review Board for Health Sciences Research at the University of Virginia approved this study. Human frontal cortex brain samples were prepared as we previously described ^3^.

### Mouse Brain Cell Extracts

Brain cortex samples were placed in a Wheaton 1 ml Dounce homogenizer containing 1 ml lysis buffer to cover the sample. Lysis buffer consisted of RIPA buffer (Bioworld, #420200242); 1% HALT™ protease inhibitor cocktail; 1% phosphatase inhibitor cocktail 2 (Sigma Aldrich, #P5726); and 1% phosphatase inhibitor cocktail 3 (Sigma Aldrich, #P0044). Brain homogenates were placed in an Eppendorf 5415C centrifuge at 10,000 rpm (8,160 x g_max_) for 10 minutes at 4° C. Finally, supernatants were transferred to new tubes and used as protein lysates for western blots.

### Atomic force and Electron microscopy for Tau proteoforms

Sample preparation and imaging were obtained exactly as described by us before ^4^.

### Human shRNA Sequences

The following plasmids were purchased from Mission shRNA library from Sigma-Aldrich: (1) NADK2.1 TRCN00000414087, (2) NADK2.2 TRCN00000424042, (3) PYCR1.1 TRCN0000038979, (4) PYCR1.2 TRCN0000038981, (5) P5CS.1 TRCN0000064849, (6) P5CS.2 TRCN0000064852, (7) NADK1 TRCN00000199808, (8) LRP1 TRCN0000230615, (9) LRP1 TRCN0000230615. These constructs are already cloned into pLKO.1, and lentivirus particles were prepared as described in a previous section. The empty backbone of the pLKO.1 was used as a control.

### Antibodies

The following antibodies were from Cell Signaling Technologies: NADK2 (Proteintech C5orf33 #26352-1-AP; 1/5000), NADK1 (cell signaling #55948; 1/1000), P5CS (Proteintech # 17719-1-AP; 1/1000), PYCR1 (Proteintech #13108-1-AP; 1/5000), PRODH (LSBio #LS-C795520; 1/1000), P5CDH (Proteintech # 11604-1-AP; 1/1000), LRP1 (cell signaling #64099; 1/1000), ColVI (Proteintech #17023-1-AP; 1/5000), T22 (Millipore Sigma #ABN454-1; 1/5000), NeuN (Cell signaling #24307; 1/1000). βIII-tubulin antibody (Tuj1; 1/10000) was a generous gift by Dr. Anthony Spano. Dr. Lester Binder provided Tau-R1 hybridoma cells (1/1000, respectively).

### Immunoblotting

Samples were resolved by SDS-PAGE using 12% acrylamide/bis-acrylamide gels and transferred to 0.22 µm nitrocellulose (Bio-Rad). Membranes were blocked with Odyssey blocking buffer (LI-COR Biosciences) and were incubated with primary antibodies and secondary IRDye-labeled antibodies diluted into antibody buffer (Odyssey blocking buffer diluted 1:1 with PBS/0.1% Tween 20). All antibody incubations were overnight at 4° C, and 3 washes of 5 minutes each with PBS/0.1% Tween 20 were performed after each antibody step. Membranes were quantitatively scanned by an Odyssey imaging station (LI-COR Biosciences).

### Other fluorescence microscopy procedures

Cells were rinsed in PBS and fixed for 15 min in 2% paraformaldehyde. Then, they were washed and permeabilized in washing buffer (0.3% Triton in PBS) once for 5 min each, washed with DPBS, and quenched with 300 mM Glycine in DPBS for 10 min. The cells were incubated for 60 min in blocking buffer (TBS-LI-COR, Biosciences #927-60001; supplemented with 1% Goat Serum, Southern Biotech #00060-01). Fixed cells were then incubated with primary (ON) and secondary (1.5 h) antibodies diluted into a blocking buffer, with several PBS washes after each antibody step. DAPI (Thermo-Scientific #62248) was incubated in the last washing step. Finally, coverslips were mounted in each well using Fluoromount-G (Thermo-Fisher #00-4958-02). Samples were imaged on a Zeiss LSM 880 inverted microscope equipped with an AiryScan 32-channel GaAsP spectral detector array, 2-PMT, a 20x/0.8 Plan Apochromat objective, a 405nm Diode laser, 488nm Argon laser, 542nm DPSS laser, and a 594/633nm HeNe laser.

### Recombinant tau

The human 2N4R tau with a C-terminal his-tag was produced by expression in BL21(DE3) Escherichia coli cells. Protein expression was induced with 0.5 mM Isopropyl β-D-1-thiogalactopyranoside (Sigma-Aldrich, I6758) for 2 hours at 37° C. Cells were pelleted by centrifugation, resuspended, and sonicated with Misonix Sonicator 3000 20 times for 30 seconds each at 60% power. Then, the E. coli lysate was centrifuged in Sorvall RC6 Plus 6 centrifuges at 18,083 x g_max_ for 25 minutes with an SLA 1500 rotor. Tau was then purified batch-wise from the supernatant using TALON Metal Affinity Resin (Takara, cat. no. 635502) according to the vendor’s instructions. Finally, purified 2N4R tau was concentrated, and the buffer was exchanged into 10 mM HEPES, pH 7.6, using Amicon Ultra-4 Centrifugal Filters (Millipore, UFC801024)^4^.

### Tau oligomers

Oligomeric tau was prepared as described earlier ^5^. The 2N4R tau isoform was adjusted to 8 µM in the oligomerization buffer: 10 mM HEPES (pH 7.6), 100 mM NaCl, 0.1 mM ethylenediaminetetraacetic acid, and 5 mM dithiothreitol (DTT). The protein was then allowed to oligomerize in the presence of 300 µM arachidonic acid (Cayman Chemicals, cat. no. 90010) for 18 hours at room temperature in the dark. Oligomerization was verified on western blots. Oligomerization buffer, prepared in parallel with oligomeric tau, containing 300 µM arachidonic acid, was used as a vehicle (control) for cell treatments ^4^. Dr. Rakez Kayed generously donated the human brain-derived tau oligomers (BDTauO).

### ATP accumulation assays

Wild-type (WT) human neurons (differentiated for 21 days) were grown in 96-well plates (black wall/clear bottom, Costar catalog number #3603). ATP was assayed using the ATPlite 1-step Luminescence Assay System, following the manufacturer’s instructions (Perkin-Elmer).

### Subcellular fractioning and Mass spectrometry protein identification

Subcellular fractioning was performed according to the vendor’s instructions from the Qproteome® Cell compartment kit (QIAGEN, Catalog #37502). 100 µg of membrane fraction from human neurons WT and NADK2-KO were sent for mass spectrometry analysis at the Biomolecules analysis core facility at UVA. Briefly, the samples were precipitated by acetone-methanol to produce a protein pellet. The pellet was re-solubilized in a digestion buffer using heat and sonication. The samples were reduced with DTT for 30 minutes at RT, then alkylated using iodoacetamide for 30 minutes at RT, followed by overnight digestion with 0.5 µg trypsin. The samples were cleaned up using beads and C18 tips. The LC-MS system consisted of a Thermo Orbitrap Exploris 480 mass spectrometer system with an Easy Spray ion source connected to a Thermo 3 µm C18 Easy Spray column (through pre-column). 10 µL of the extract was injected, and the peptides eluted from the column by an acetonitrile/0.1 M acetic acid gradient at a flow rate of 0.3 µL/min over 2 hours. The nanospray ion source was operated at 1.9 kV. The digest was analyzed using the rapid switching capability of the instrument, acquiring a full scan mass spectrum to determine peptide molecular weights followed by product ion spectra (Top10 HCD) to determine the amino acid sequence in sequential scans. The data was analyzed by database searching using the Sequest search algorithm against Uniprot Human. The peptides and proteins identified for the sample were analyzed in Scaffold 5.3.3 software.

### NADK2, P5CS, and PYCR1 Gene knockout

The LentiCRISPRv2 system (Addgene #98290) was used to assess CRISPR-Cas9-mediated gene knockout of the human NADK2 in progenitors ReNCell MV cell line. Two human guide RNA sequences were chosen from the CRISPR pick portal (https://portals.broadinstitute.org/gppx/crispick/public). Primer design, ligation, and transformation were followed from the LentiCRISPRv2 and lentiGuide-Puro: lentiviral CRISPR/Cas9 and single guide RNA. Then, plasmids were packed into lentiviral particles (as described before) to transduce cells. Human ReNCell MV cell lines were plated in 96-well plates in a confluence of ∼100 cells per well. After 24 h, cells were transduced with the NADK2 LentiCRISPR-virus. After 48 h of transduction, complete antibiotic selection (1 µg/ml puromycin) was applied to the genetically modified cells before proceeding to experiments. After 4 weeks, the resistant cells were detached and expanded. After the expansion, a second round of selection was conducted. Then, the knockout human ReNCell MV cells were differentiated for 21 days for each experiment. The efficiency of gene knockout was established by WB before and after the differentiation process. sgRNA sequences used in this study were sgNADK2-1; TCCTAGGAATGAGGGAATTG, and sgNADK2-2; TGGCATAAACCCTGTACCTG. sgP5CS-1; GAAGCATGCCAAGAGAATCG and sgP5CS-2; TGCCAATGGAACCCACCCAA. sgPYCR1-1; GAAGTTGACACCCCACAACA and sgPyCR1-2; CGTGCCTGTGGCATACACGG.

### Measurement of intracellular proline levels

A previously described protocol ^6–8^ was adapted as follows. Briefly, cells were harvested and lysed in cold PBS with 1% Triton X-100. The cells were centrifugated to remove cellular debris (10,000 x g). The supernatants were transferred to a new Eppendorf and boiled in a water bath. Intracellular amino acids were extracted by boiling for 10 min. After a centrifugation step (5 min, 4°C, 15,000 x g), the supernatant was free of proteins, and intracellular proline was determined as described below. Briefly, 100 µL of the supernatant was incubated with 100 µL of acid-ninhydrin (0.25 g ninhydrin dissolved in 60/40 glacial acetic acid/6 M phosphoric acid) and 100 µL of glacial acetic acid for 1 h at 100°C. The reaction was stopped, and samples were incubated on ice for 5 min, and the mixture was extracted with 200 µL toluene. Carefully, the toluene phase was separated, and absorbance at 520 nm was used to determine the proline concentration. All the reactions were performed in triplicates, and a standard curve ranging in concentrations of 0.05–1 mM proline was paralleled to determine the proline concentration of samples ^6–8^.

### Biotin labeling of tau protein and endocytosis assay

A previously described protocol was adapted to label oligomeric tau ^9,10^. EZ-Link Sulfo-NHS-Biotin (2 mg, Thermo-Scientific #A39258) was reconstituted in water to 2 mM. TauOs were incubated with the biotin at 1:1 molar ratio (total tau:biotin) for 30 minutes at room temperature or 2 hours on ice. To remove unbound biotin and enable buffer exchange into PBS, the solution was centrifuged four times at 10,000 ×g for 15 minutes each in Amicon Ultra-0.5 Centrifugal Filters (Millipore, UFC501096), with the addition of PBS to the solution remaining above the filter at each step. Human neurons were then exposed to 300 nM biotinylated RTauO (bt-RTauO). After 1, 3, and 6 hours of incubation, bt-RTauO that remained bound to the outer surface of the plasma membrane were reduced and removed with 50 mM DTT for 1 hour at 37°C. Cells were washed three times with PBS for 15 minutes and lysed under native conditions with hypotonic buffer (10 mM Tris HCl, pH 7.4; 1.5 mM MgCl2; 10 mM KCl). Protein samples were mixed with 1X NuPage LDS Sample Buffer (Invitrogen #NP0007) without reducing agents and analyzed by western blotting using Streptavidin-IRDye800CW (LI-COR #926-32230) to detect biotinylated tau that had been endocytosed.

### Reduced Nicotinamide adenine dinucleotide (NADH) and Reduced Nicotinamide adenine dinucleotide phosphate (NADPH) measurements and in vivo 2P-FLIM imaging

#### Human neurons 2P-FLIM imaging and processing

Human neuronal cultures were grown in 35-mm glass-bottom dishes, maintained at 37°C in 5% CO2/95% air on the stage of the Zeiss LSM-980 NLO microscope. The laser was tuned to 740 nm with an average power of 7 mW at the specimen plane, and NAD(P)H fluorescence was collected using a 450 to 500 nm emission filter (Objective lens, 40 ×). For each experiment, 10 fields of view were recorded in the descanned mode, and then each field of view was subjected to a 40-second acquisition in the non-descanned mode. The laser power and acquisition time were selected to ensure enough photons per pixel while avoiding photodamage to cells.

FLIM images were processed with SPCImage software (v8.6; Becker & Hickl) at maximum-likelihood fitting at 3-component lifetimes for human neurons. Based on Blacker et al^11^, enzyme-bound NADH and NADPH lifetimes fall into specific Ꚍ2 (1400-4400 picoseconds) and Ꚍ3 (>4400 picoseconds); their fractions of a_2_% and a_3_% therefore express the relative abundance of the enzyme-bound coenzymes, charted here. All FLIM image parameter data were exported for further processing in Fiji software (https://hpc.nih.gov/apps/Fiji.html). Here, from thresholded, normalized photon images, pixel-ROIs are isolated based on the more prominent intensity of the mitochondrial morphology, which were applied to all FLIM parameter data, creating a mitochondria-specific data pool. Those results were analyzed using a custom code in Python software, where all parameters were first examined and filtered for obvious outliers (insignificant numbers). The FLIM metrics of interest – NADH-a_2_% and NAD(P)H-a_3_% were extracted as indicators for changes in the mitochondrial metabolic OXPHOS state and plotted for publication-ready charts and statistics in Microsoft Excel. Surgical preparation of animals, *in vivo* 2P-FLIM imaging of live mouse cortex, and Imaging analysis were performed exactly as described by us before ^1–3^

### Statistics

The paired t-test was used to analyze all 2P-FLIM assays. Since thousands of ROIs were obtained per image, the average ROI for each field of view was calculated to reduce the sample size and, thus, the number of false positives. All data were assumed to be normally distributed. A p-value of less than 0.05 was considered statistically significant.

---

**Extended DATA #1:**
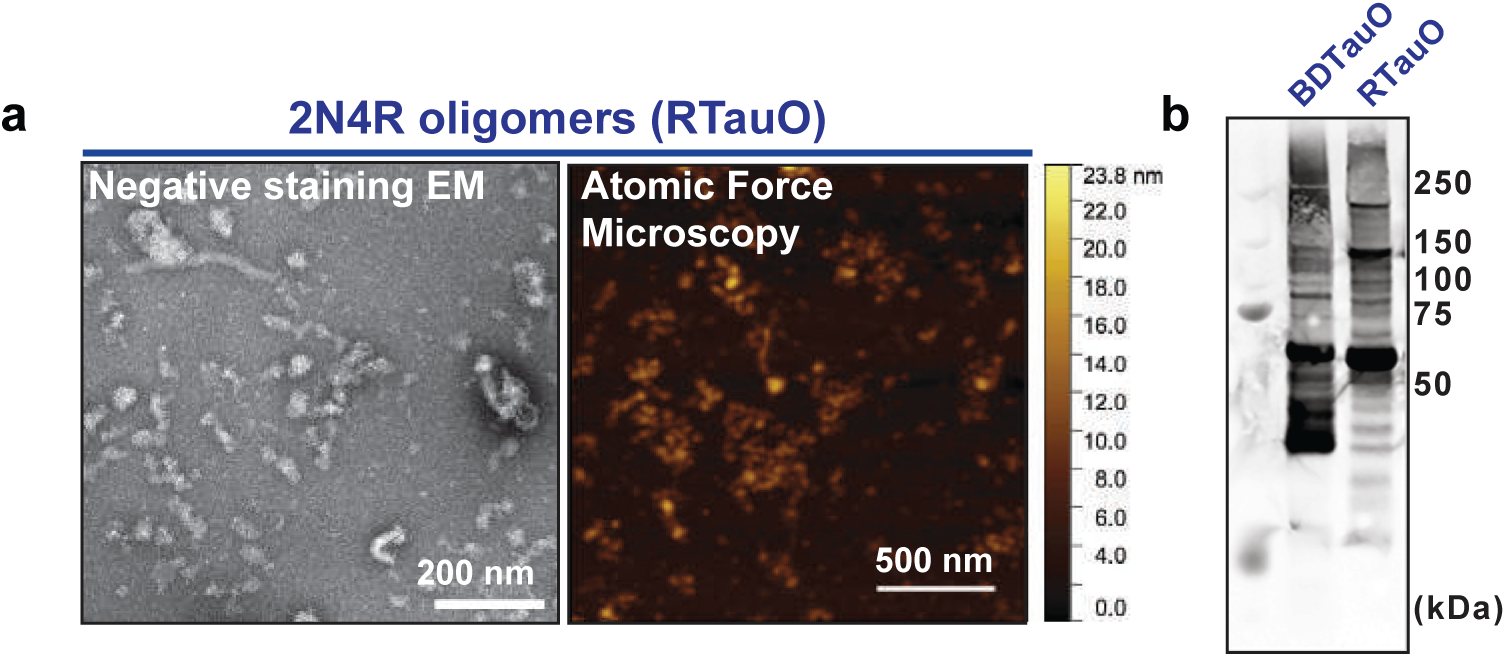
Recombinant his-tagged 2N4R human tau and human brain-derived tau oligomers used in this study. a. Human 2N4R tau oligomeric visualized by negative staining electron microscopy (left image) and visualized by atomic force microscope (right image). b. Comparison of human brain-derived tau oligomers (BDTauO) and human recombinant tau oligomers (RTauO) by WB using Tau5 antibody. Dr. Rakez Kayed generously gifted human brain-derived Tau oligomers.

**Extended DATA #2:**
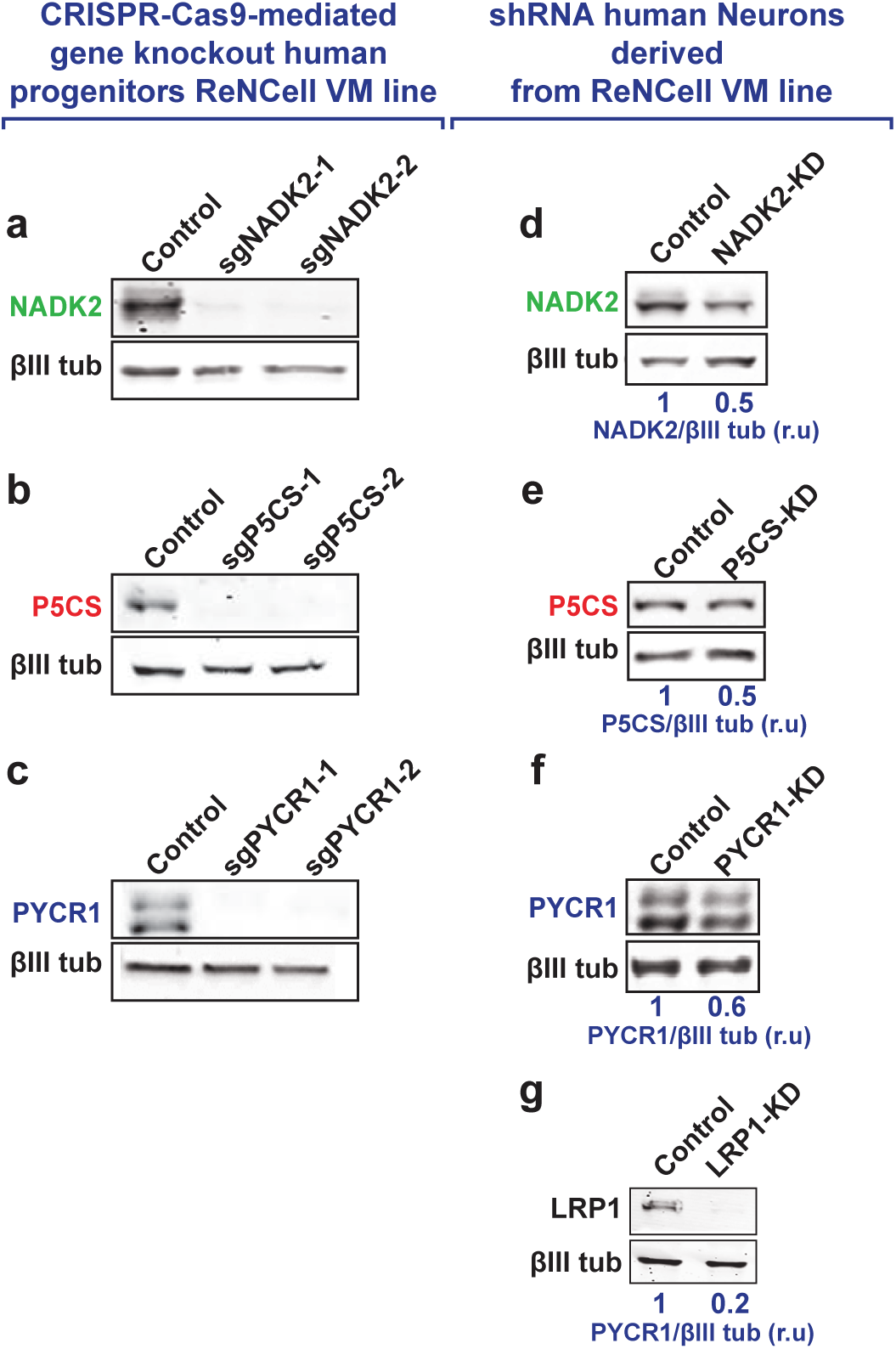
Biochemical validation of NADK2, P5CS, and PYCR1 knockout and knockdown in human neurons. (a) NADK2, (b) P5CS, (c) PYCR1 knockout neurons were made using CRISPR/Cas9 technology. Neurons were harvested, and the respective knockout efficiency was measured by WB using the indicated antibodies and normalized to the expression level of βIII tubulin contained in the same samples. (d) NADK2, (e) P5CS, and (f) PYCR1 expression were down-regulated using lentiviral-mediated delivery of shRNA sequences. Cells were harvested, and the corresponding knockdown efficiency was measured by WB using the indicated antibodies and normalized to the expression level of βIII tubulin contained in the same samples.

**Extended DATA #3:**
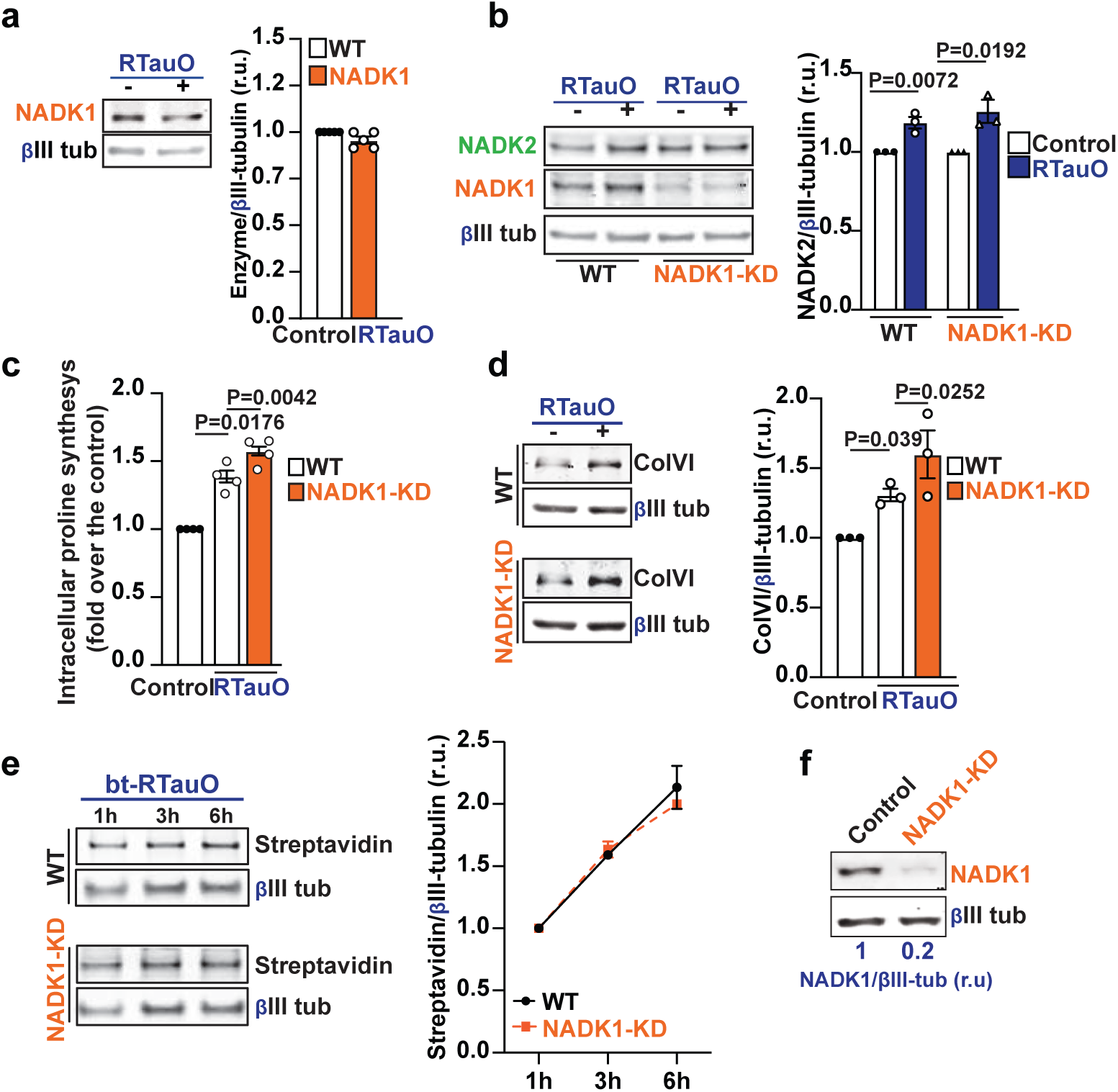
TauO-mediated effects on the mitochondrial NADK2 are independent of NADK1 expression. **a.** NADK1 expression was assessed by WB in cell extracts from WT human neurons treated with either vehicle (control) or 100 nM of RTauO for 24 hours and normalized to the expression level of bIII tubulin contained in the same samples. Statistical analyses were performed using a two-tailed unpaired student’s t-test. (n=5). Error bars represent +/-s.e.m. **b.** WT and NADK1 knocked down human neurons were treated with vehicle (control) or 100 nM of RTauO for 24 hours. NADK2 and NADK1 levels were measured by WB and normalized to the expression level of bIII tubulin contained in the same samples. Statistical analyses were performed using a two-tailed unpaired student’s t-test. (n=3). Error bars represent +/- s.e.m. Intracellular proline levels (**c**; n=4) and ColVI synthesis (**d**; n=4) were monitored in WT or NADK1 knocked-down neurons treated with RTauO for 24 hours, as described before. **e.** An endocytosis assay of bt-RTauO in WT and NADK1 knocked-down neurons was performed as described in Fig. 2f-g. Line graphs show the quantification obtained using 3 independent experiments and analyzed using a two-tailed unpaired student’s t-test. **f.** Lentiviral-mediated delivery of shRNA sequences against NADK1 (NADK1-KD) efficiency was ∼80%.

**Extended DATA #4:**
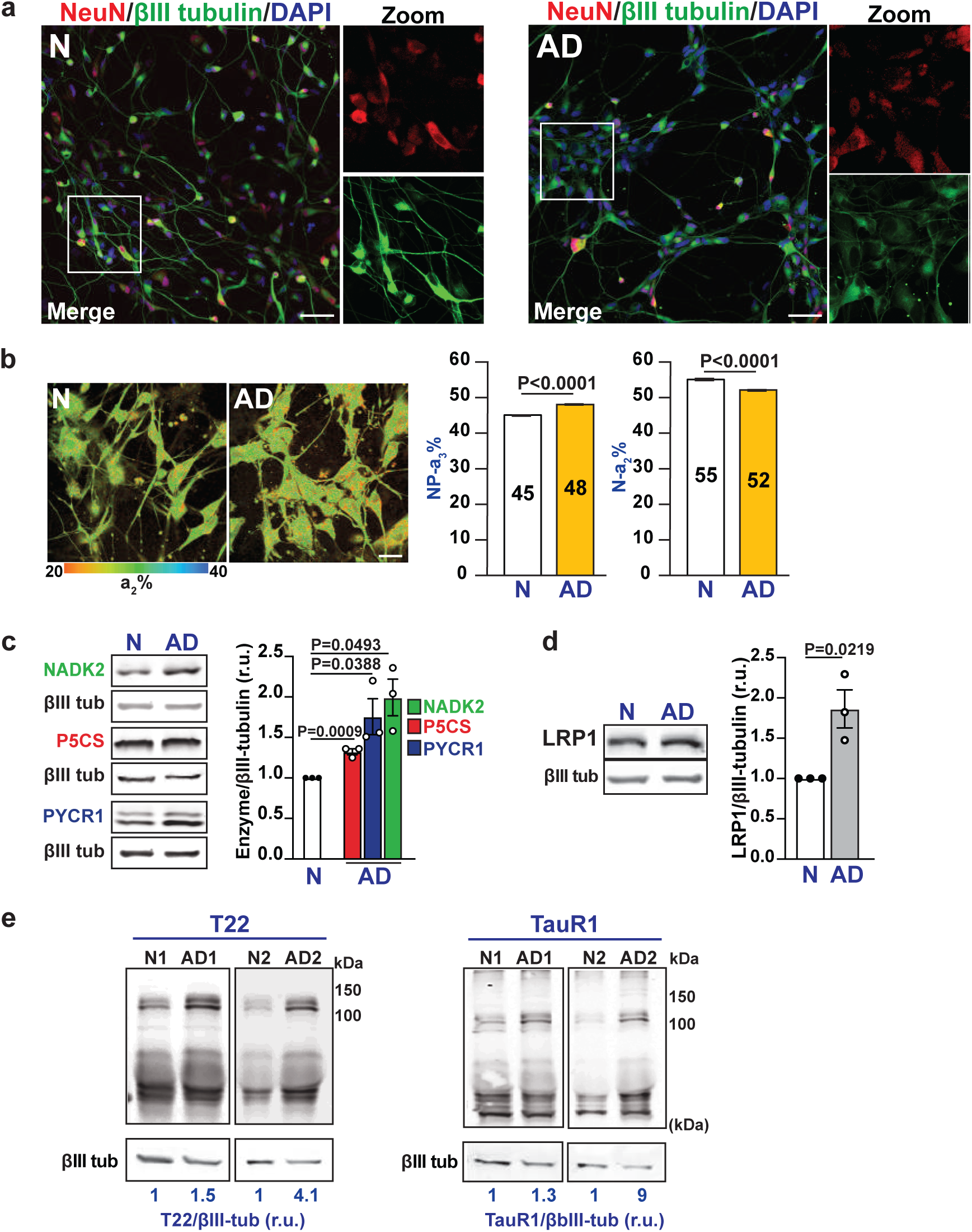
Human AD iNeurons exhibit increased NADK2, NADPH, and the proline synthesis pathway along with an increment of tau proteoforms. **a.** Expression of neuronal markers 3 weeks post induction iNeurons. Representative immunofluorescence from age and sex-matched cognitively normal control (N) and Alzheimer’s disease donor (AD) iNeurons using NeuN (red), βIII tubulin (green), and DAPI (blue). Scale bars, 50 μm. **b.** Representative 2P-FLIM images of human iNeurons, obtained by direct conversion of dermal fibroblasts from cognitively normal control (patient #6; male, 76 years old) or AD (patient #4; male, 76 years old) donors. The experiment was performed with iNeurons from 3 AD and 3 cognitively normal donors. Scale bars, 10 μm. Statistical analyses were performed using a two-tailed unpaired student’s t-test. Error bars represent +/- s.e.m. **c.** NADK2, P5CS, and PYCR1 expression levels in healthy and AD iNeurons were analyzed by WB against the indicated proteins. **d.** LRP1 expression was analyzed by WB of protein samples obtained from either 3 healthy control or 3 AD iNeurons. Statistical analyses in c and d were performed using a two-tailed unpaired student’s t-test. Error bars represent +/- s.e.m. (n=3) **f.** WB analyzed the expression of high molecular weight tau species detected with T22 antibody-positive tau proteoforms and TauR1 antibody-positive for human tau.

